## Supplemental Figures 1-7 for "Maintenance of neuronal TDP-43 expression requires axonal lysosome transport"

**Supplemental figures and legends**

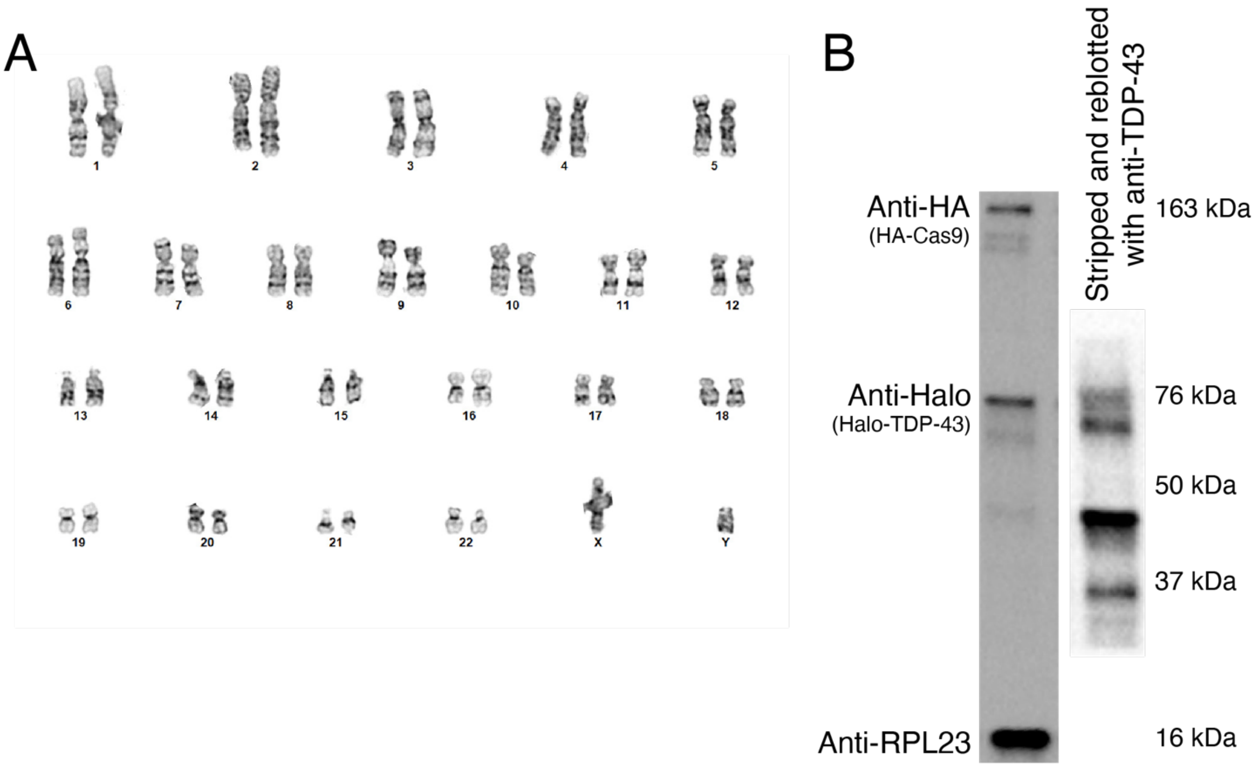


Figure S1 (related to Figure 1): i11w-hT iPSCs have a normal karyotype.

A) Karyotype of Halo-TDP-43 iPSCs shows a normal karyotype (44 chromosomes + XY), demonstrating that CRISPR editing did not cause any chromosomal abnormalities.

B) Left blot shows Cas9, Halo-TDP-43, and RPL23 staining (Anti-HA, Anti-Halo, and Anti-RPL23, respectively), demonstrating TDP-43 is HaloTagged. On right, blot was stripped and re-probed with anti-TDP-43, showing both HaloTagged TDP-43 and untagged TDP-43.

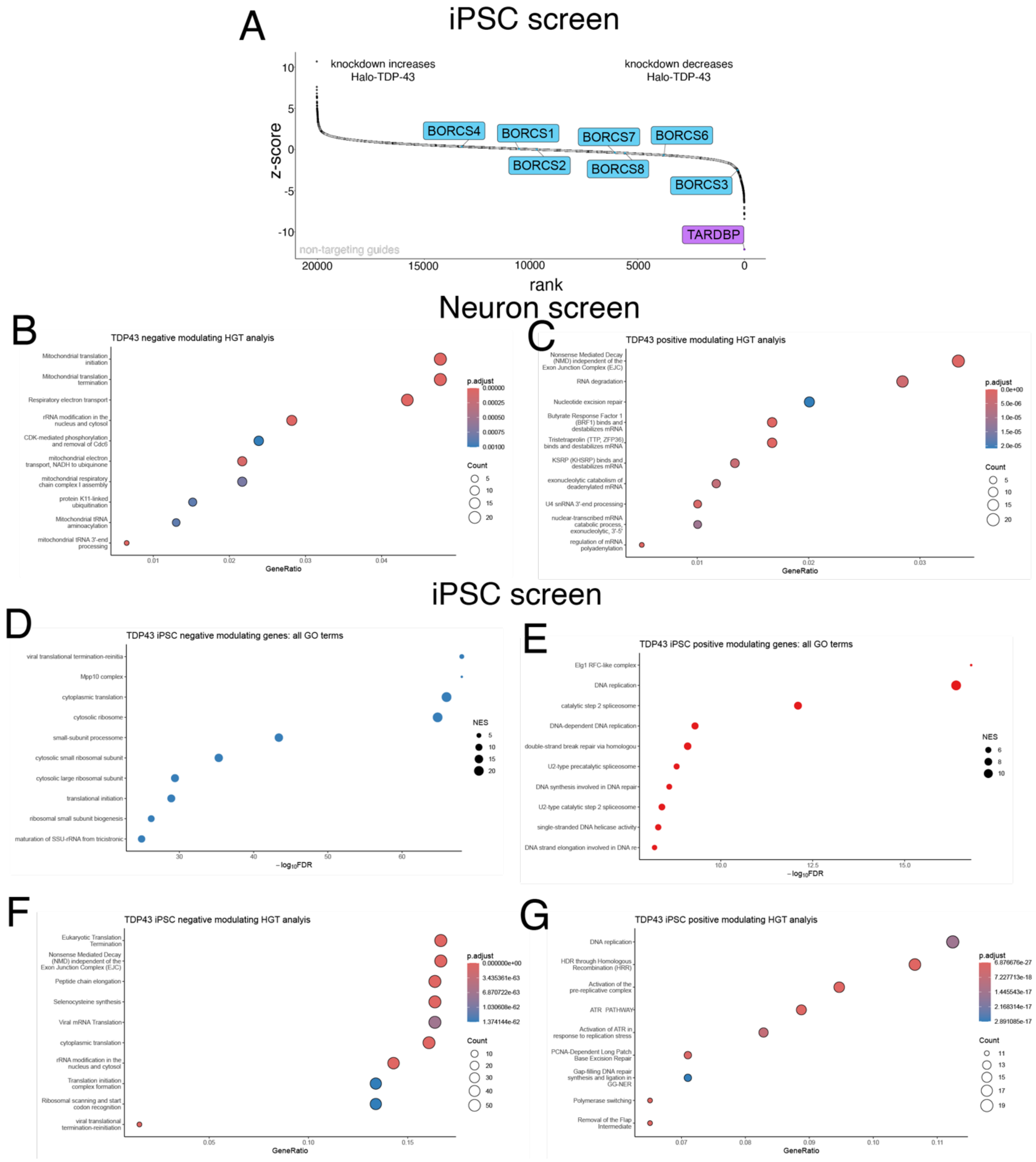


Figure S2 (related to Figure 2): iPSC screen shows different hits than neuron screen.

A) Rank plot of iPSC screen results showing genes whose KD increases Halo-TDP-43 levels (left) and genes whose KD decreases Halo-TDP-43 levels, including TDP-43 itself (right). BORC genes are indicated in blue. Non-targeting (control) guides indicated in gray.

B) HGT analysis of screen hits that decrease Halo-TDP-43 levels in neurons also shows many mitochondrial terms, as was seen in the GO analysis.

C) HGT analysis of screen hits that increase Halo-TDP-43 levels in neurons shows many RNA metabolism hits.

D) GO analysis of hits that decrease Halo-TDP-43 levels in iPSCs shows many ribosome and translation-related terms, a striking difference from the hits found in neurons. The FDR is from the calculated permutation p value of 1000 iterations.
E) GO analysis of hits that increase Halo-TDP-43 levels in iPSCs shows DNA synthesis, spliceosome, and DNA repair terms. The FDR is from the calculated permutation p value of 1000 iterations.
F) HGT analysis of hits that decrease Halo-TDP-43 levels in iPSCs shows translation, ribosome, and NMD-related terms. The FDR is from the calculated permutation p value of 1000 iterations.
G) HGT analysis of hits that increase Halo-TDP-43 levels in iPSCs shows DNA related terms. The FDR is from the calculated permutation p value of 1000 iterations.

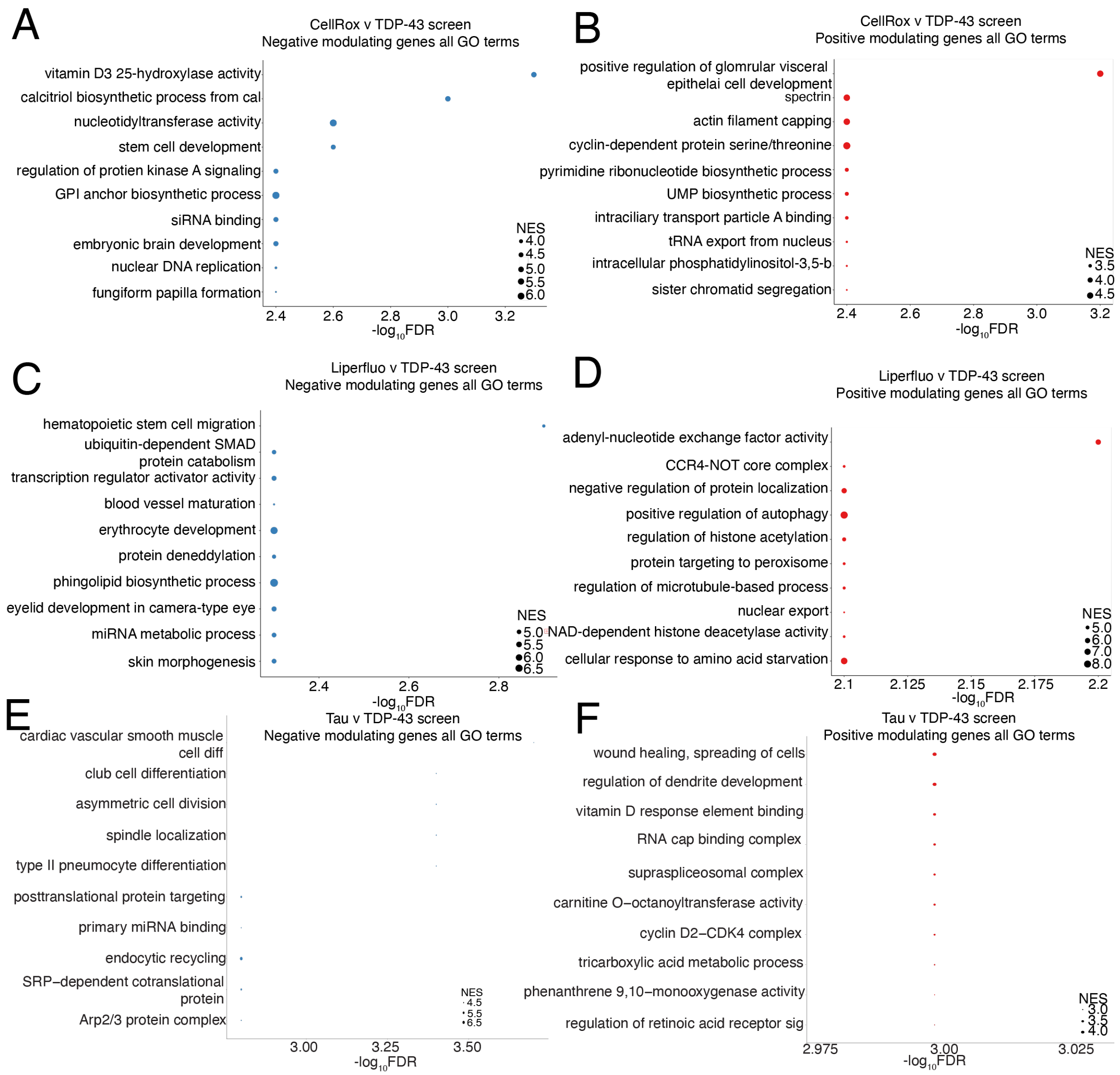


Figure S3 (related to Figure 3): Meta-analyses of Halo-TDP-43 and published CRISPRi FACS screens.

A) GO analysis of genes that negatively modulate readouts for CellRox and Halo-TDP-43 screens show DNA and signaling related terms. The FDR is from the calculated permutation p value of 1000 iterations.
B) GO analysis of genes that positively modulate readouts for CellRox and Halo-TDP-43 screens show translation related terms. The FDR is from the calculated permutation p value of 1000 iterations.
C) GO analysis of genes that negatively modulate readouts for Liperfluo and Halo-TDP-43 screens show terms related to ubiquitination, neddylation, and lipids. The FDR is from the calculated permutation p value of 1000 iterations.
D) GO analysis of genes that positively modulate readouts for Liperfluo and Halo-TDP-43 screens show adenosine, acetylation, and autophagy related terms. The FDR is from the calculated permutation p value of 1000 iterations.
E) GO analysis of genes that negatively modulate readouts for Tau and Halo-TDP-43 screens show terms related to ubiquitination, neddylation, and lipids. The FDR is from the calculated permutation p value of 1000 iterations.
F) GO analysis of genes that positively modulate readouts for Tau and Halo-TDP-43 screens show adenosine, acetylation, and autophagy related terms. The FDR is from the calculated permutation p value of 1000 iterations.

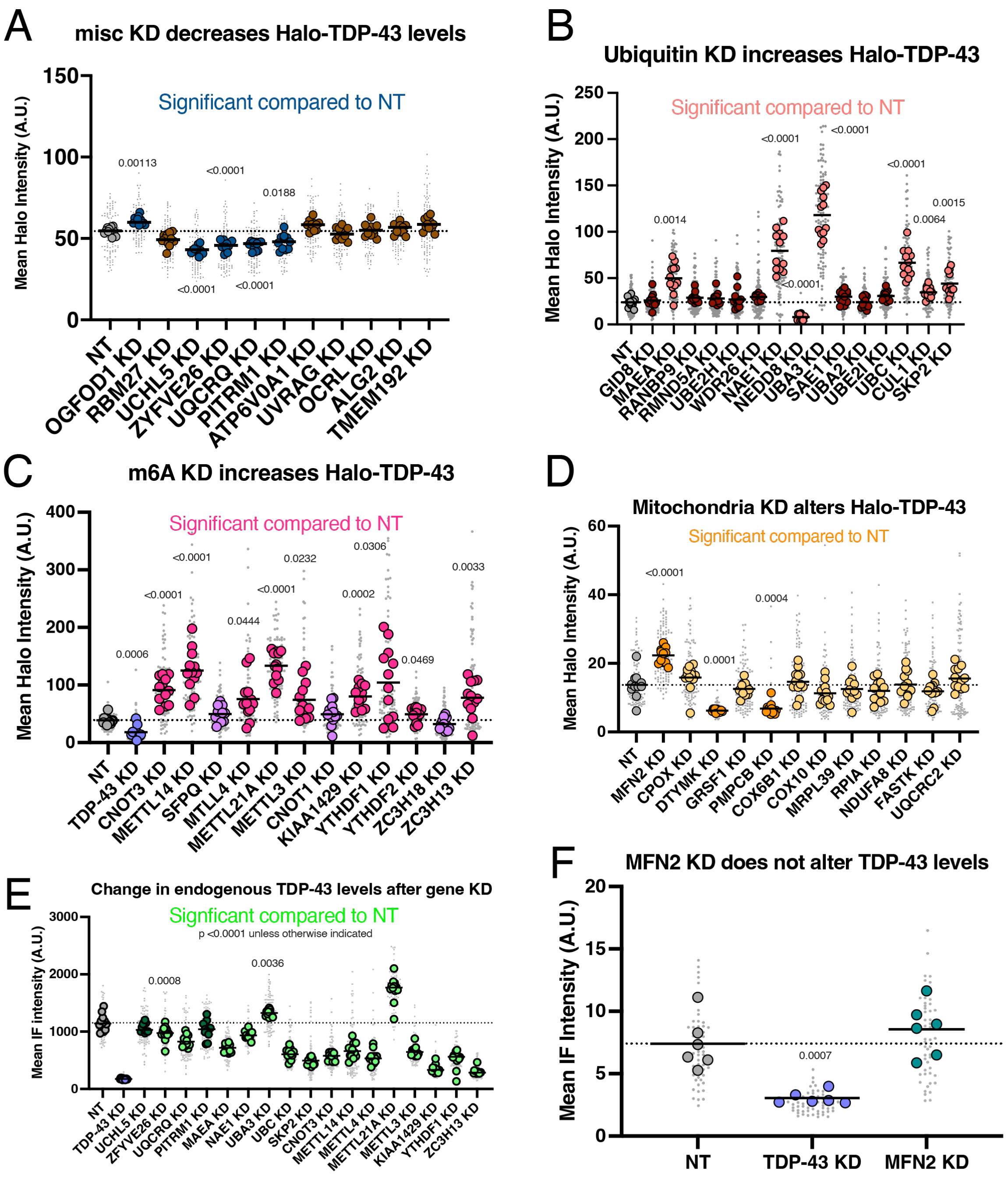


Figure S4 (related to Figure 4): Validation of neuron screen results show many genes alter TDP-43 levels.

A) Quantification of miscellaneous KD Halo-TDP-43 microscopy. OGFOD1 KD increases Halo-TDP-43 levels while UCHL5 KD, ZYFVE26 KD, UQCRQ KD, and PITRM1 KD decrease Halo-TDP-43 levels compared to a non-targeting (NT) guide, indicated by dark blue dots. N=12 wells per genotype, 9 images per well (small grey dots). Significant p-values indicated on graph.

B) Quantification of ubiquitin-associated gene KD Halo-TDP-43 microscopy. MAEA, NAE1, UBA3, UBC, CUL1, and SKP2 KD all increase Halo-TDP-43 levels compared to a non-targeting (NT) guide, indicated by orange dots. NEDD8 KD decreases Halo-TDP-43 levels due to a strong survival phenotype. N=12 wells per genotype, 9 images per well (small grey dots). Significant p-values indicated on graph.

C) Quantification of m6A-associated gene KD Halo-TDP-43 microscopy. CNOT3, METTL14, MTLL4, METTL21A, METTL3, KIAA1429, YTHDF2, and ZC3H13 KD all increase Halo-TDP-43 levels compared to a non-targeting (NT) guide, indicated by magenta dots. N=12 wells per genotype, 9 images per well (small grey dots). Significant p-values indicated on graph.

D) Quantification of mitochondria-associated gene KD Halo-TDP-43 microscopy. MFN2 KD increases Halo-TDP-43 levels while DTYMK and PMPCB KD decrease Halo-TDP-43 levels compared to a non-targeting (NT) guide, indicated by dark blue dots. N=12 wells per genotype, 9 images per well (small grey dots). Significant p-values indicated on graph.

E) Quantification of TDP-43 immunofluorescence (IF) microscopy. TDP-43 KD significantly decreases TDP-43 levels by IF. ZFYVE26, UQCRQ, , MAEA, NAE1, UBC, SKP2, CNOT3, METTL14, METTL4, METTL3, KIAA1429, YTHDF1, and ZC3H13 KD all decrease TDP-43 IF levels (although not all decrease Halo-TDP-43 levels), while UBA3 and METTL21A KD increase TDP-43 IF levels, indicated by lime dots. N=12 wells per genotype, 9 images per well (small grey dots). Significant p-values indicated on graph.

F) Quantification of MFN2 KD TDP-43 immunofluorescence microscopy. MFN2 KD does not significantly change untagged TDP-43 levels compared to a NT guide, indicated by teal dots, while TDP-43 KD does significantly decrease TDP-43 IF levels. N=6 wells per genotype, 9 images per well (small grey dots). Significant p-values indicated on graph.

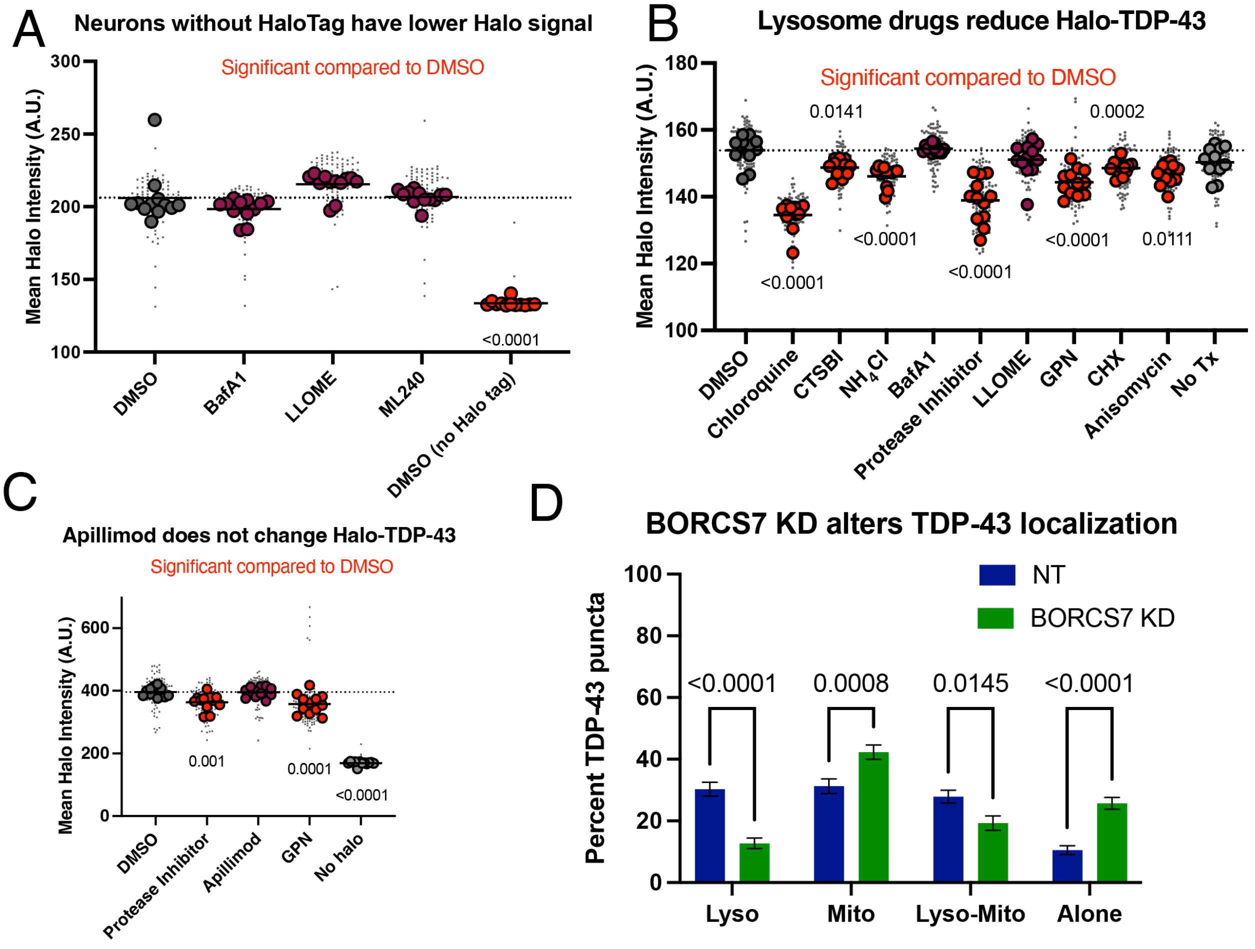

Figure S5 (related to Figure 6): Lysosome inhibitors alter Halo-TDP-43 levels.

A) Additional drug treatments from experiment shown in Figure 5B that do not change Halo-TDP-43 levels. An untagged control line (“no Halo tag”) shows significantly less Halo signal than the DMSO control (repeated from Figure 5B).
B) Replication of drug treatment experiment shows the same trend as previous experiment.
C) Apillimod does not alter levels of Halo-TDP-43.
D) Quantification of Halo-TDP-43-organelle association after BORCS7 KD. Halo-TDP-43 is significantly reduced on lysosomes, with a small increase of Halo-TDP-43 on mitochondria. More Halo-TDP-43 is transported not on either lysosomes or mitochondria in BORCS7 KD neurons.

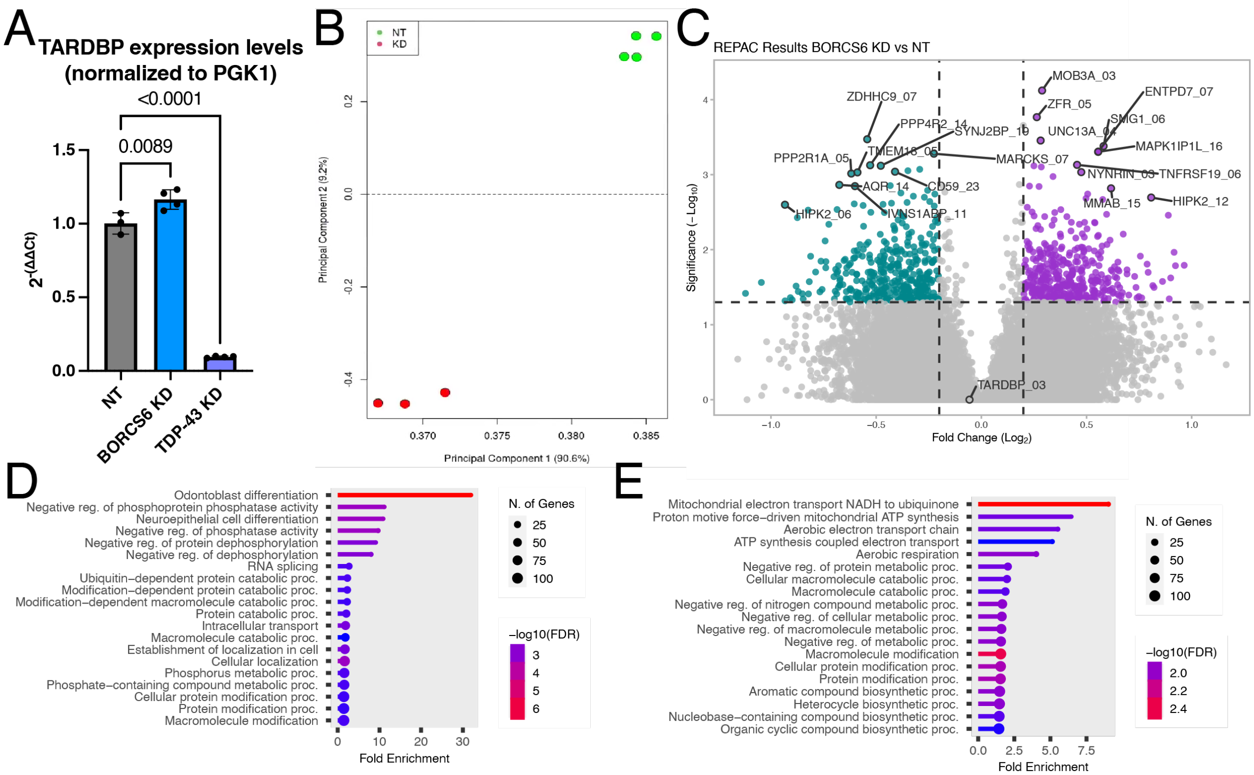


Figure S6 (related to Figure 7): TDP-43 mRNA levels are unchanged after BORCS6 KD.

A) qPCR of NT, BORCS6 KD, and TDP-43 KD neurons showing decreased TDP-43 mRNA levels in TDP-43 KD, and slight, but significant increase in TDP-43 mRNA in BORCS6 KD. Normalized to PGK1. NT n=3, BORCS6 KD n=4, and TDP-43 KD n=4.
B) Principal component analysis plot of RNA sequencing data shows most of the variance between the samples (90%) can be explained by a single principal component and that the samples separate based on knockdown status.
C) REPAC results comparing BORCS6 KD to NT show changes in poly adenylation across the transcriptome, with increased usage of an alternative polyA site compared to the reference site on the right of the volcano, and decreased use of the polyA site compared to the reference site on the left side. TDP-43 (TARDBP) polyA is not changed in BORC KD neurons.
D) For alternative polyA sites that result in a lengthening event (right side of volcano in C), GO BP shows enrichment in pathways including splicing, intracellular transport, and posttranslational modifications.
E) For alternative polyA sites that result in a shortening event (left side of volcano in C), GO BP shows enrichment in terms related to catabolism, metabolism, and biosynthesis.

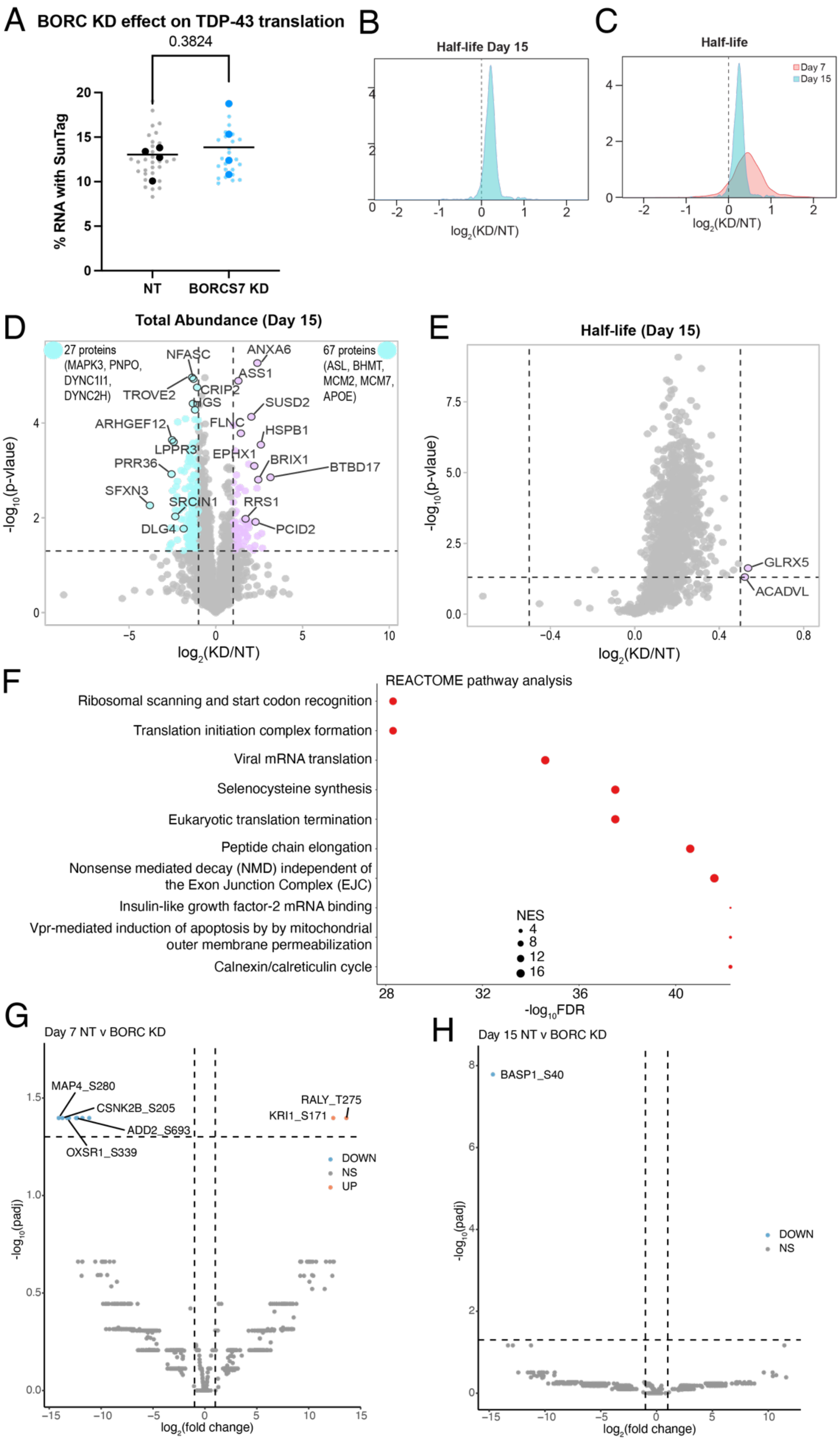
A
BORC KD effect on TDP-43 translation
0.3824
% RNA with SunTag
NT
BORCS7 KD
B
Half-life Day 15
log2(KD/NT)
C
Half-life
Day 7
Day 15
log2(KD/NT)
D
Total Abundance (Day 15)
27 proteins (MAPK3, PNPO, DYNC1I1, DYNC2H)
67 proteins (ASL, BHMT, MCM2, MCM7, APOE)
NFASC
TROVE2
CRIP2
HGS
ARHGEF12
LPPR3
PRR36
SFXN3
SRCIN1
DLG4
ANXA6
ASS1
SUSD2
FLNC
HSPB1
EPHX1
BRIX1
BTBD17
RRS1
PCID2
-log10(p-vlaue)
log2(KD/NT)
E
Half-life (Day 15)
GLRX5
ACADVL
-log10(p-vlaue)
log2(KD/NT)
F
REACTOME pathway analysis
Ribosomal scanning and start codon recognition
Translation initiation complex formation
Viral mRNA translation
Selenocysteine synthesis
Eukaryotic translation termination
Peptide chain elongation
Nonsense mediated decay (NMD) independent of the Exon Junction Complex (EJC)
Insulin-like growth factor-2 mRNA binding
Vpr-mediated induction of apoptosis by by mitochondrial outer membrane permeabilization
Calnexin/calreticulin cycle
NES
4
8
12
16
-log10FDR
G
Day 7 NT v BORC KD
MAP4_S280
CSNK2B_S205
ADD2_S693
OXSR1_S339
RALY_T275
KRI1_S171
DOWN
NS
UP
-log10(padj)
log2(fold change)
H
Day 15 NT v BORC KD
BASP1_S40
DOWN
NS
-log10(padj)
log2(fold change)

Figure S7 (Related to figure 8): Protein turnover is increased at d15 in BORC KD neurons.

A) TDP-43 translation is not altered upon BORCS7 KD, as measured by fraction of RNA puncta with SunTag puncta by microscopy. N=4 wells, 8 images quantified per well.
B) Density plot of total protein half-life shows longer protein half-lives in BORCS6 KD neurons compared to NT neurons at d15.
C) Overlay of protein half-life density plots at d7 (red) and d15 (teal) show both days have longer protein half-lives in BORCS6 KD neurons.
D) Volcano plot of total protein abundance comparing BORCS6 KD to a non-targeting guide at d15. Some proteins (purple) have increased abundance, while a larger number (teal) have decreased abundance. Non-significant or low $\log_2$(fold change) proteins are indicated in grey. Vertical lines at $\log_2$(fold change) of 1 and -1, horizontal line at p-value = 0.05.
E) Volcano plot of total protein half-life at d15 show many proteins have longer half-lives ($\log_2$(fold change) > 0) in BORCS6 KD neurons as compared to neurons expressing a non-targeting guide. Non-significant or low $\log_2$(fold change) protein turnovers are indicated in grey. Horizontal line at p=0.05, vertical lines at $\log_2$(fold change) of 0.5 or -0.5.
F) Gene ontology enrichment analysis using proteins with longer half-lives in BORCS6 KD neurons shows REACTOME pathways related to ribosomes, translation, and mRNA binding.

**Supplementary table**

Table S1: MAGeCKFlute robust ranked algorithm (RRA) results for genes that increased Halo-TDP-43 levels in i3Neurons.

Table S2: MAGeCKFlute robust ranked algorithm (RRA) results for genes that decreased Halo-TDP-43 levels in i3Neurons.

Table S3: MAGeCKFlute robust ranked algorithm (RRA) results for genes that increased Halo-TDP-43 levels in iPSCs.

Table S4: MAGeCKFlute robust ranked algorithm (RRA) results for genes that decreased Halo-TDP-43 levels in iPSCs.

Table S5: Individual guide sequences cloned for screen validation. Secondary screen results summaries and figures used in are indicated in last three columns.
